# Heterologous vaccination regimens with self-amplifying RNA and Adenoviral COVID vaccines induce robust immune responses in mice

**DOI:** 10.1101/2021.01.28.428665

**Authors:** Alexandra J Spencer, Paul F McKay, Sandra Belij-Rammerstorfer, Marta Ulaszewska, Cameron D Bissett, Kai Hu, Karnyart Samnuan, Anna K. Blakney, Daniel Wright, Hannah R Sharpe, Ciaran Gilbride, Adam Truby, Elizabeth R Allen, Sarah C Gilbert, Robin J Shattock, Teresa Lambe

## Abstract

Several vaccines have demonstrated efficacy against SARS-CoV-2 mediated disease, yet there is limited data on the immune response induced by heterologous vaccination regimens using alternate vaccine modalities. Here, we present a detailed description of the immune response, in mice, following vaccination with a self-amplifying RNA (saRNA) vaccine and an adenoviral vectored vaccine (ChAdOx1 nCoV-19/AZD1222) against SARS-CoV-2. We demonstrate that antibody responses are higher in two dose heterologous vaccination regimens than single dose regimens. Neutralising titres after heterologous prime-boost were at least comparable or higher than the titres measured after homologous prime boost vaccination with viral vectors. Importantly, the cellular immune response after a heterologous regimen is dominated by cytotoxic T cells and Th1^+^ CD4 T cells which is superior to the response induced in homologous vaccination regimens in mice. These results underpin the need for clinical trials to investigate the immunogenicity of heterologous regimens with alternate vaccine technologies.

## Introduction

A number of vaccines against SARS-CoV-2 have reached late-stage clinical trials with encouraging efficacy readouts and mass vaccination schemes already initiated across several countries. Multiple vaccine technologies are being advanced ^1^, but there is limited information on how these vaccine modalities may work in combination. With ongoing clinical trial assessment of numerous SARS-CoV-2 vaccines and mass vaccination incentives, there is a recognised real-world scenario where individuals may be vaccinated with different vaccine modalities. However, the utility of vaccination regimens combining different types of vaccine approaches remains to be determined.

Self-amplifying RNA (saRNA) typically encodes the alphaviral replicase and a target antigen. Upon entry into the cytoplasm, the RNA is amplified with subsequent translation of the target antigen ^2^. The SARS-CoV-2 saRNA vaccine has been demonstrated to be highly immunogenic in preclinical animal models ^3^ and is progressing through clinical trial assessment with promising results. Adenoviruses are a frequently used viral vector vaccine technology, they can be rapidly made to GMP at large scale, and a single vaccination can be sufficient to provide rapid immunity in individuals ^4^. In particular, chimpanzee derived adenoviruses (ChAd) have good safety profiles, whilst inducing strong cellular and humoral immune response against multiple target disease antigens ^5–7^. Previous studies have demonstrated efficacy after vaccination with ChAdOx1 nCoV-19/AZD1222 against SARS-CoV-2 mediated disease ^8^, with concomitant high titre humoral immune responses and a measurable Th1-dominated cellular immune response ^9,10^. Importantly, an immune profile consistently observed across different animal species ^11,12^.

In this study, the immunogenicity of saRNA and ChAdOx1 vaccines expressing full-length SARS-CoV-2 spike protein were assessed in mice following vaccination with different combinations of vaccine modalities. We demonstrate robust antibody responses following heterologous vaccination regimens, with high titre neutralising antibodies. The cellular immune response is dominated by cytotoxic T cells secreting IFNγ and TNFα and antigen specific CD4+ T cells of a Th1 phenotype, with significantly higher antigen specific responses observed following heterologous vaccination than those responses induced in single vaccine regimens. These results underpin the need for clinical trial assessment of immunisation regimens with alternate vaccine modalities.

## Results

### Heterologous vaccination potentiates SARS-CoV-2 spike-specific antibody response

It was previously shown that homologous prime-boost immunization with saRNA ^3^ or ChAd 11 induced a SARS-CoV-2 spike-specific IgG response with neutralisation capacity. In this series of experiments, we compared the immune response induced by heterologous, homologous and single dose vaccination with ChAd and saRNA vaccine modalities. Heterologous vaccination with either ChAd or saRNA prime and alternative boost (i.e. ChAd-saRNA or saRNA-ChAd) or two doses of saRNA induced the highest IgG responses, compared to mice vaccinated with a single dose of either ChAd or saRNA (Fig. 1A and Fig. S1A), with strong correlation between the two independent IgG ELISA methods (Fig. S1B). This IgG response showed mixed profile, with mainly IgG2 (IgG2a, IgG2b in both mouse strains and IgG2c in CD1 mice) and IgG1 subclasses (in both mouse strains) (Fig. 2A and B), which was similar to previously reported results after vaccination with ChAd alone ^13^ or saRNA alone ^3^. Comparable amounts of SARS-CoV-2 spike-specific IgM were detected in all vaccinated groups (Fig. 1B), while heterologous vaccination with saRNA-ChAd induced higher serum SARS-CoV-2 spike-specific IgA levels compared to single vaccination with either vaccine, in both CD1 and BALB/c mice (Fig. 1B).

**Fig 1:**
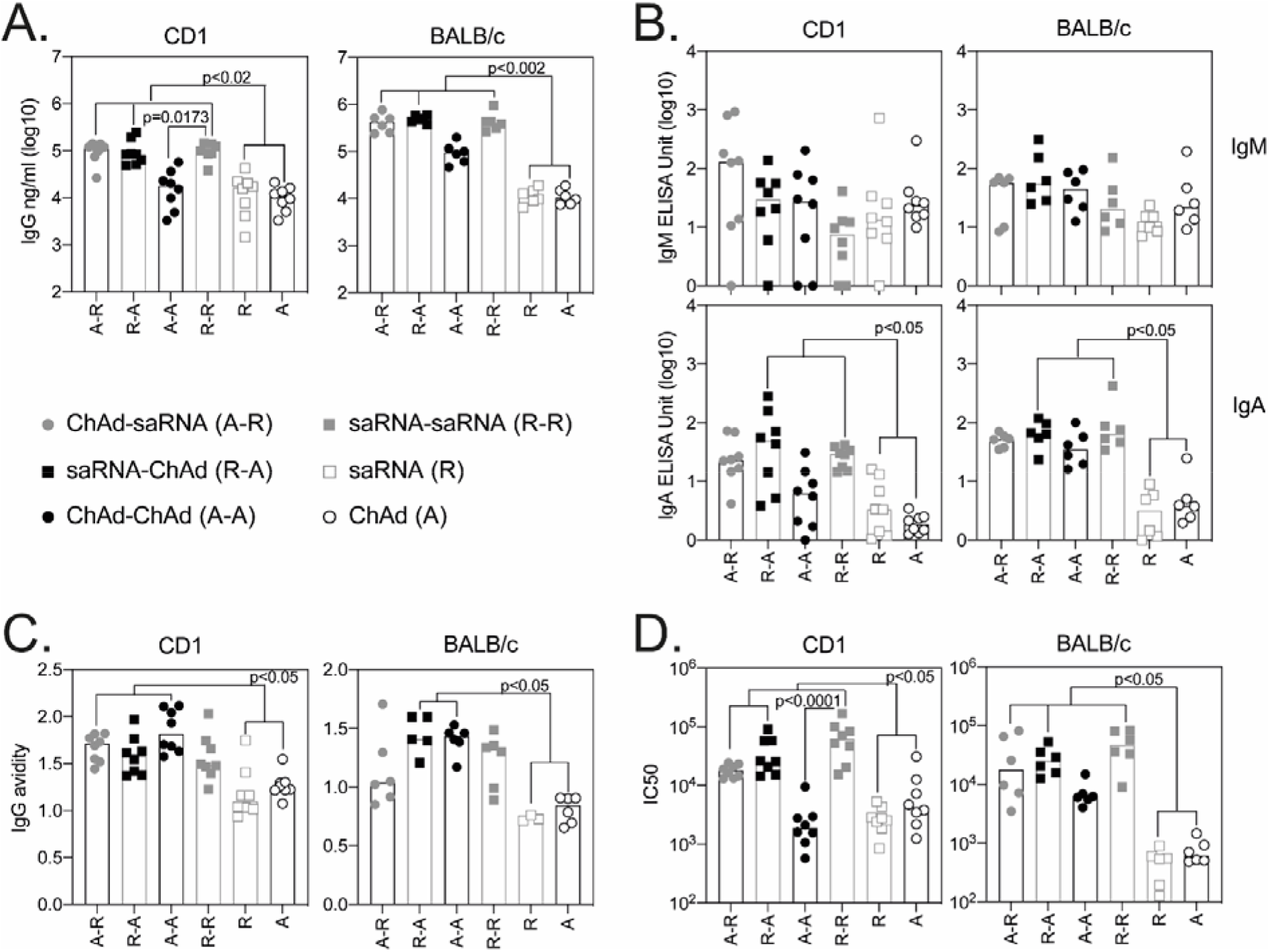
Antibody responses following ChAd and saRNA vaccination. Antibody responses were measured in the serum of CD1 (n=8) and BALB/c (n=6) mice collected 3 weeks after the final immunisation. Graphs show SARS CoV-2 spike-specific IgG **(A.)**, IgM and IgA **(B.)** and IgG avidity **(C.)** measured by ELISA, and SARS-CoV-2 pseudotyped virus neutralisation (IC50) **(D.)**. Individual mice are represented by a single data point, bars represent the median response in each group (CD1 n=8; BALB/c n=6). Data in each graph was analysed with a Kruskal-Wallis and post-hoc positive test to compare differences between vaccination groups, p values indicate significant differences between groups.

**Fig 2:**
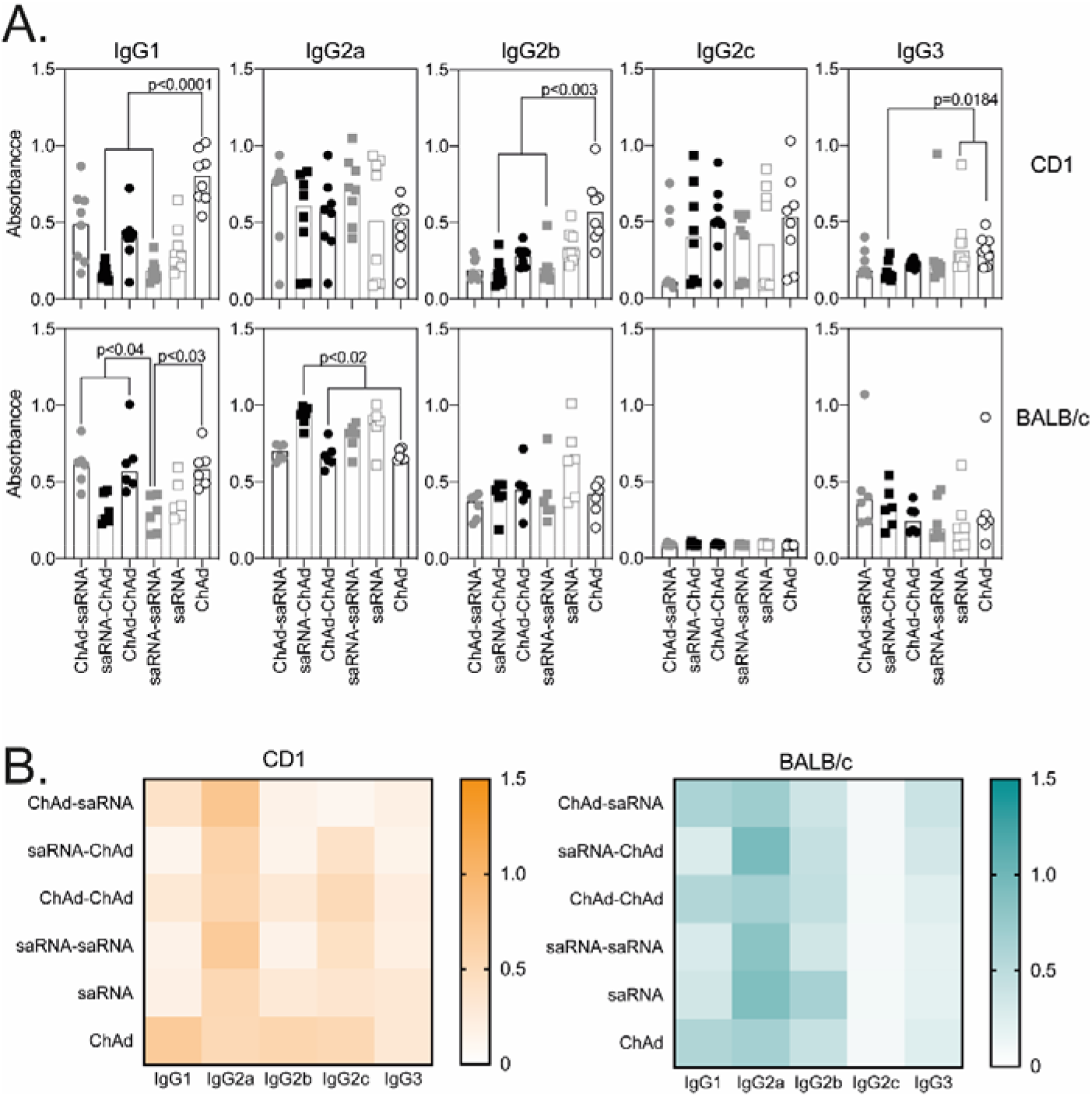
SARS-CoV-2 spike-specific IgG subclasses following ChAd and saRNA vaccination. For detection of IgG subclasses, each sample was diluted to 1 IgG ELISA Unit. Graphs show optical density measured against each IgG subclass where individual data points were expressed as an OD and shown here as scatter dot plots with bars showing the median **(A.)**, followed by the heatmap summary representation with median response in each group to each IgG subclass **(B.)**. Individual mice are represented by a single data point, bars represent the median response in each group (CD1 n=8; BALB/c n=6) with serum collected 3 weeks after the final vaccination. Data in each graph was analysed with a Kruskal-Wallis and post-hoc positive test to compare differences between vaccination groups, p-values indicate significant differences between groups.

Two doses of vaccine were also shown to increase the avidity of the IgG towards SARS-CoV-2 spike (Fig. 1C) compared to single administration of either vaccine, while antibody-mediated neutralisation was measured across all groups of vaccinated mice, with heterologous and two dose saRNA inducing significantly higher levels of neutralisation compared to single dose vaccination regimens (Fig. 1D). Not surprisingly, a correlation was measured between neutralisation and levels of IgG /IgA SARS-CoV-2 spike-specific antibodies (Fig. S1C).

Flow cytometry staining enabled identification of SARS-CoV-2 spike-specific B cells in the spleen of BALB/c mice. The data demonstrated that all vaccination regimens induced a similar number of antigen specific B cells (Fig. S2B), with similar numbers of germinal centre (GC) and isotype class switched B cells observed in all groups of vaccinated mice. Formation of GC is important for generation of long-lived memory B cells and the data demonstrates that changing vaccine modalities did not impact formation of GC B cells.

### Heterologous vaccination induces strong Th1 type response

T-cell mediated immunity was also investigated following heterologous or homologous ChAdOx1 and saRNA vaccination regimens. The highest IFNγ response detected by ELISpot was observed in mice that received a heterologous combination of vaccines, this increase was only statistically significant when compared to single administration of saRNA in both strains of mice (Fig. 3A and B), or saRNA-ChAd compared to ChAd in BALB/c mice (Fig. 3B). In agreement with earlier reports, ELISpot responses were primarily directed towards the S1 portion of the spike protein (Fig S3A), with a consistent breadth of response measured with all vaccine combinations and in both strains of mice (Fig. 3A and B).

**Fig 3:**
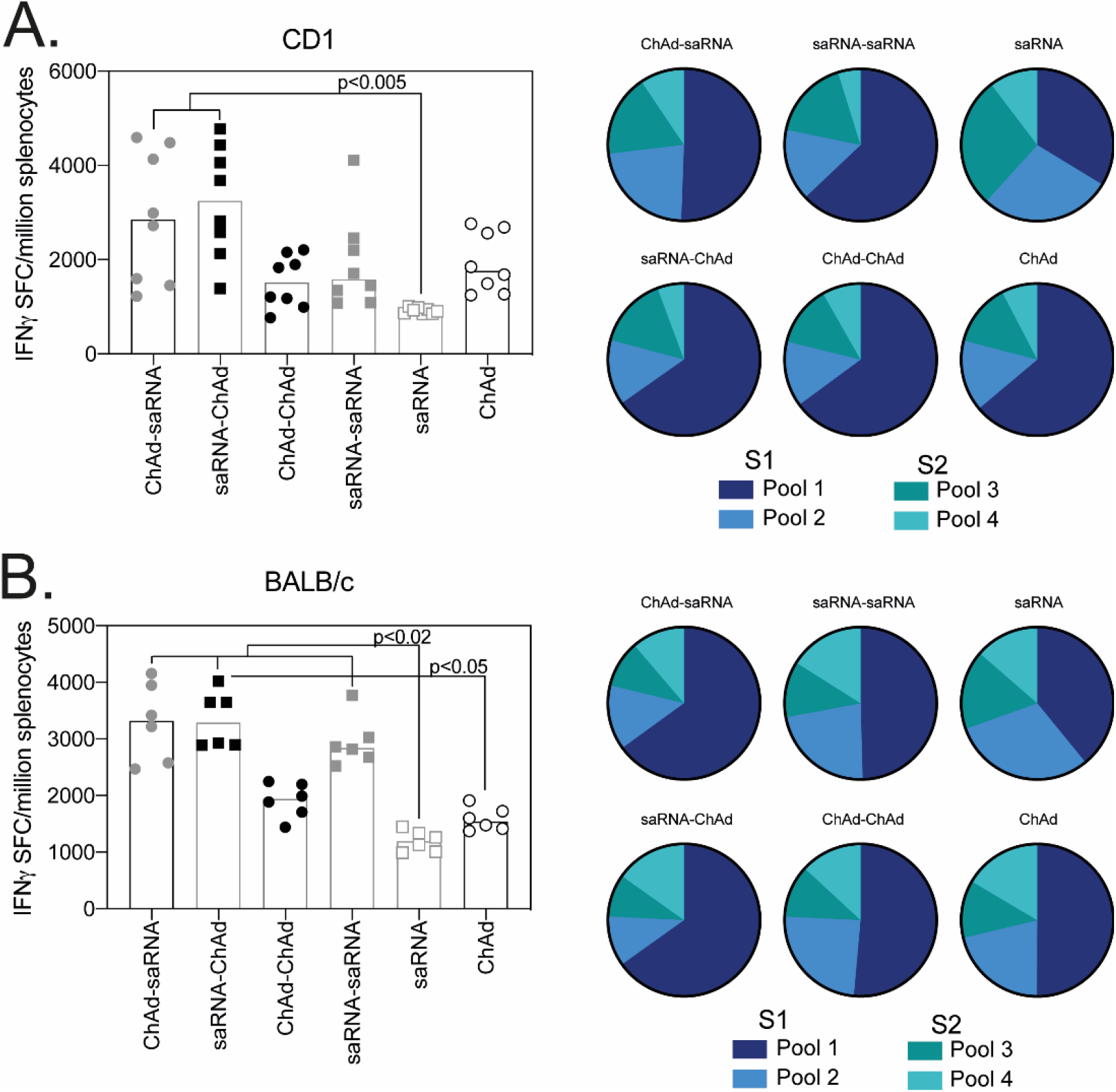
Breadth of T cell response measured by ELISpot. Graphs represent the total spike specific IFNγ response (sum of peptide pools) measured in outbred CD1 (n=8) **(A.)** or inbred BALB/c (n=6) **(B.)** 3 weeks after the final vaccination. Pie charts represent the response to each peptide pool as a proportion of total response. Data points represent individual mice and bars represent the median response in each group. Data in each graph was analysed with a Friedman test and post-hoc Dunn’s multiple comparison to compare between vaccination regimens, p values are indicated on the graph.

Phenotype and functional capacity of the T cell response was measured by intracellular cytokine staining, together with memory T cell marker staining. Consistent with previously published data, saRNA and ChAd vectors induced antigen specific CD4^+^ T cells of a Th1 bias, with minimal IL4 and IL10 production measured and a response dominated by production of IFNγ and TNFα, regardless of the vaccination regimen (Fig. 4A). Antigen specific CD4^+^ T cells displayed mixed T effector (Teff) and T effector memory (Tem) phenotype, with no statistically significant differences observed in the total number of cells (Fig. 4A) and cell subsets (Fig. S3C). As before, in mice the cell-mediated response following vaccination was dominated by CD8^+^ T cells, with much higher levels of IFNγ and TNFα (in addition to upregulation of CD107a) seen in all vaccine groups (Fig. 4B) compared to CD4^+^ T cell responses. In addition, the highest number of antigen specific T cells, in both outbred and inbred strains of mice, was measured following heterologous vaccination, regardless of vaccine order (Fig. 4B) with antigen specific CD8^+^ T cells displayed a predominantly Teff phenotype.

**Fig 4:**
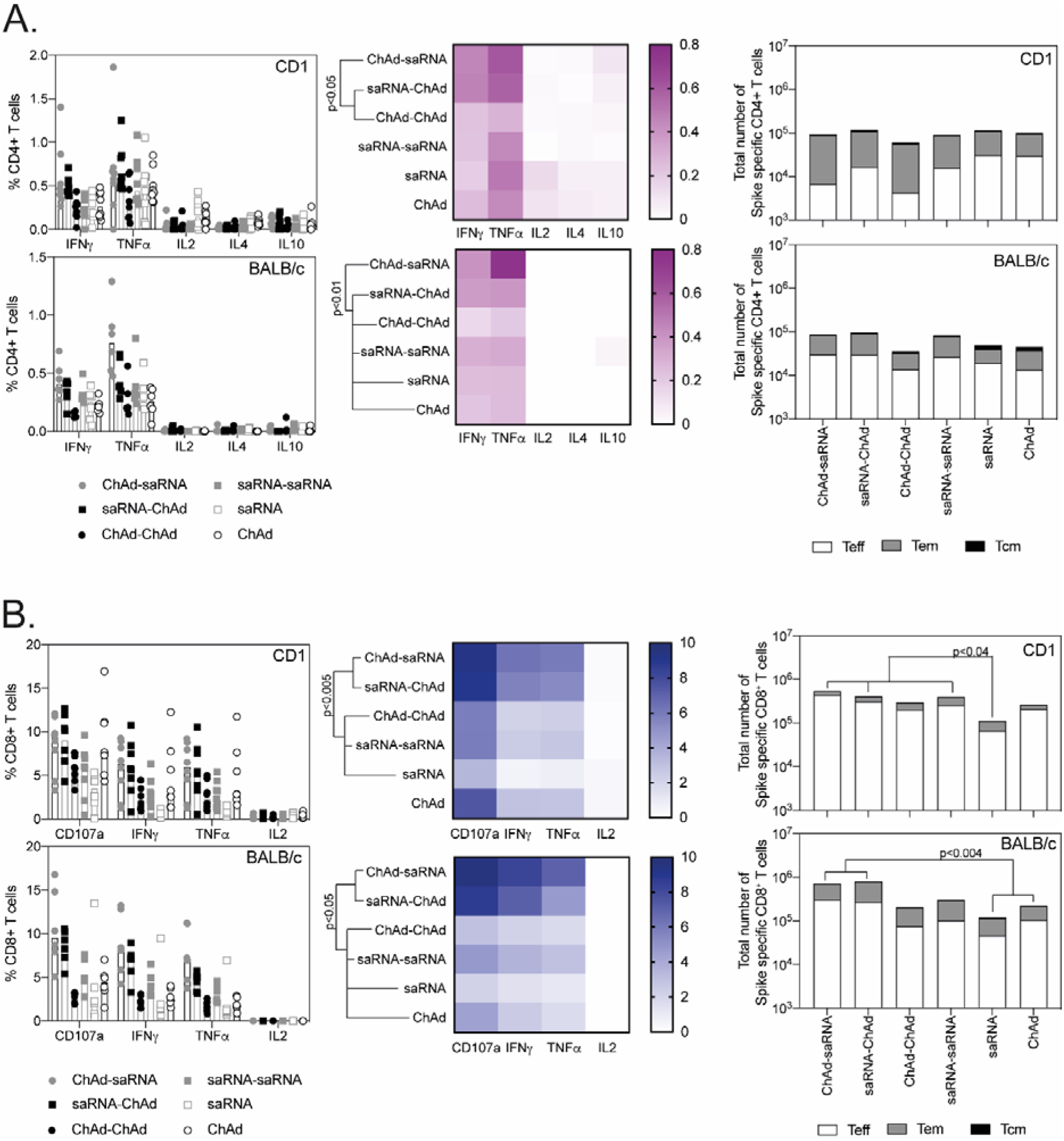
Phenotype of the T cells response following vaccination. CD1 (n=8) and BALB/c (n=6) splenocytes harvested 3 weeks after the final vaccination, were stimulated for 6 hours with pools of overlapping SARS-CoV-2 peptides prior to staining for effector and memory T cells markers and intracellular cytokines. **A.)** Graphs show the frequency of the spike-specific CD4^+^ T cells responses (left) in CD1 (top panel) and BALB/c mice (bottom panel), heatmaps (middle) show the proportion of the response producing each cytokine and total number of antigen specific cells of a T effector (Teff), T effector memory (Tem) or T central memory (Tcm) phenotype (right). **B.)** Graphs show the frequency of the spike-specific CD8^+^ T cells responses (left) in CD1 (top panel) and BALB/c mice (bottom panel), heatmaps (middle) show the proportion of the response producing each cytokine and total number of antigen specific cells of a T effector (Teff), T effector memory (Tem) or T central memory (Tcm) phenotype (right). Data points indicate individual mice, bars represent the median response in each group. Total numbers of each population are displayed in Fig S3C. Data in each graph was analysed with a two-way anova comparing the effect of vaccination regimen and cytokine production or T cell phenotype, followed by a post-hoc Tukey’s multiple comparison test to compare between vaccination regimens, p values showing overall differences between vaccination groups are indicated on the graph.

## Discussion

The current pandemic and extraordinary efforts to develop effective vaccines with subsequent mass vaccination roll-out has highlighted the ‘real-world’ practicalities of global vaccination campaigns. There are approximately 20 vaccines in Phase 3 clinical trial assessment, and several vaccines have already reported efficacy with subsequent emergency licensure granted in some countries. While each individual vaccine candidate has been thoroughly tested for safety and efficacy, there have been no studies reported to date that have examined the safety, efficacy or any added benefit of mixed modality vaccinations. Given the real-world vaccination initiatives that are being progressed, there are scenarios wherein an individual receives a vaccine prime and a boost dose from different manufacturers or of different vaccine types. This current pre-clinical study examined the cellular and humoral immune responses in mice following vaccination with either the ChAdOx adenoviral vector or the saRNA LNP in homologous or heterologous prime-boost combinations and strongly supports the need for clinical trial assessment of heterologous prime-boost regimens.

All prime-boost regimens elicited high levels of SARS-CoV-2 spike-specific antibodies with neutralisation capacity and high avidity, levels that were greater than single vaccines alone. Heterologous vaccination regimens induced some of the highest antibody responses post-vaccination, with neutralising titres after heterologous prime-boost at least comparable to or higher than those achieved after homologous vaccination with ChAdOx1 nCoV-19. While homologous saRNA and ChAd induced higher antibody responses than single dose regimens, which is consistent with previous data in mice, pigs and NHPs ^11,12^. We have demonstrated in NHPs, that homologous vaccination with ChAdOx1 nCoV-19 results in protection against disease, with more recent data, in hamsters, demonstrating that a single immunisation with ChAdOx1 nCoV-19 protects against disease induced by the variants of concern B.1.351 and B.1.1.7 (van Doremalen et al submitted). Importantly in human clinical trials, strong enhancement of the antibody response was observed following a booster dose of ChAdOx1^14^, with this regimen shown to be efficacious against SARS-CoV-2 disease in late-stage clinical trials ^8^. While recent real word data, in elderly frail people, has demonstrated vaccine effectiveness after the first dose of ChAdOx1nCoV-19 at 80.4% (95% CI 36.4-94.5) with broadly similar effectiveness measured after RNA (Pfizer) vaccination (Hymas et al, Lancet preprint).

While there are no defined correlates of protection from clinical trials, Rhesus Macaques studies have demonstrated a clear role for neutralising antibodies and also CD8^+^ T cells in protecting against disease^15^. In agreement, human studies have demonstrated neutralising antibodies and T cells play an important role in preventing severe disease and augmenting recovery from COVID-19^16^. Both vaccine modalities also elicited high numbers of antigen specific T cells, which were further increased in the heterologous regimens. The majority of the IFNγ ELISpot response was directed against the S1 spike protein, particularly the first half (317 AA) which does not include the RBD. The cell-mediated response was dominated by cytotoxic T cells, with heterologous regimens inducing higher frequencies and total numbers of antigen specific CD8^+^ T cells. Although CD4^+^ T cell responses were overall lower in frequency and number than CD8^+^ T cell responses, all vaccination regimens elicited CD4^+^ T cell responses of a Th1 type response, even in the Th2 biased BALB/c background, a response indicating that the potential for antibody dependent enhancement (ADE) and subsequent enhancement of respiratory disease (ERD) caused by Th2-type lung immunopathology is reduced ^17–19^.

Vaccination regimens that induce a broad immune response (humoral and cell-mediated) will likely be the best option for long-term protection against COVID-19. It remains to be determined if the higher antibody titres following heterologous vaccination regimens, as measured here, results in longer lived immunity with a broader humoral response. Critically, the logistical challenges of administering vaccines in a rapidly evolving landscape of mass vaccination schemes, coupled with limited global supply, underpins the need to generate data on mixing vaccine modalities. It will be important to clinically assess if the mixed modality regimens have an altered or lessened reactogenicity profile, and most importantly the ability to augment protection against disease or onward transmission. Importantly some of these questions will be addressed in a recent clinical study recruiting up to 820 participants to receive combinations of two different SARS CoV-2 vaccines which has been initiated in the UK (https://comcovstudy.org.uk/home). The data described herein reinforces the need for this and other clinical trials to assess the safety, immunogenicity and efficacy of heterologous vaccination regimens.

## Methods

### Ethics Statement

Mice were used in accordance with the UK Animals (Scientific Procedures) Act under project license number P9804B4F1 granted by the UK Home Office. Age matched animals were purchased from commercial suppliers as a batch for each experiment and randomly split into groups on arrival at our facility. Animals were group housed in IVCs under SPF conditions, with constant temperature and humidity with lighting on a 13:11 light-dark cycle (7am to 8pm). For induction of short-term anaesthesia, animals were anaesthetised using vaporised IsoFlo^®^. All animals were humanely sacrificed at the end of each experiment by an approved Schedule 1 method.

### Animals and Immunizations

Outbred CD1Hsd:ICR (CD-1) (Envigo) (n=8 per group), Crl:CD1 (ICR) (Charles River) (n=8 per group) and inbred BALB/cOlaHsd (BALB/c) (Envigo) (n=6 per group) mice of 7 weeks of age, were immunized intramuscularly (i.m.) in the musculus tibialis with 10^8^ infectious units (iu) of ChAdOx1 nCoV-19 ^12^, 1μg saRNA ^3^ or received no prime vaccination. Mice were boosted (or primed for mice receiving only a single vaccination) i.m. with the relevant vaccine candidate 4 weeks later. All mice were sacrificed 3 weeks after the final vaccination with serum and spleens collected for analysis of humoral and cell-mediated immunity.

### Pseudotype virus neutralisation assay

A HIV-pseudotyped luciferase-reporter based system was used to assess the neutralization ability of sera from vaccinated mice. In brief, CoV S-pseudotyped viruses were produced by co-transfection of 293T/17 cells with a HIV-1 gag-pol plasmid (pCMV-Δ8.91, a kind gift from Prof. Julian Ma, St George’s University of London), a firefly luciferase reporter plasmid (pCSFLW, a kind gift from Prof. Julian Ma, St George’s University of London) and a plasmid encoding the S protein of interest (pSARS-CoV2-S) at a ratio of 1:1.5:1. Virus-containing medium was clarified by centrifugation and filtered through a 0.45 μm membrane 72 h after transfection, and subsequently aliquoted and stored at −80 °C. For the neutralization assay, heat-inactivated sera were first serially diluted and incubated with virus for 1 h, and then the serum-virus mixture was transferred into wells pre-seeded Caco2 cells. After 48 h, cells were lysed, and luciferase activity was measured using Bright-Glo Luciferase Assay System (Promega). The IC50 neutralization was then calculated using GraphPad Prism (version 8.4). Statistical analyses were performed on log transformed data.

### Antigen-specific IgG ELISA

The antigen-specific IgG titres in mouse sera were assessed by a semi-quantative ELISA. MaxiSorp high binding ELISA plates (Nunc) were coated with 100 μL per well of 1 μg mL−1 recombinant SARS-CoV-2 protein with the pre-fusion stabilized conformation in PBS. After overnight incubation at 4 °C, the plates were washed 4 times with PBS-Tween 20 0.05% (v/v) and blocked for 1 h at 37 °C with 200 μL per well blocking buffer (1% BSA (w/v) in PBS-Tween-20 0.05%(v/ v)). The plates were then washed and the diluted samples or a 5-fold dilution series of the standard IgG added using 50 μL per well volume. Plates were incubated for 1 h at 37 °C, then washed and secondary antibody added at 1:2000 dilution in blocking buffer (100 μL per well) and incubated for 1 hr at 37 °C. After incubation and washes, plates were developed using 50 μL per well SureBlue TMB (3,3⍰, 5,5⍰-tetramethylbenzidine) substrate and the reaction stopped after 5 min with 50 μL per well stop solution (Insight Biotechnologies). The absorbance was read on a Versamax Spectrophotometer at 450 nm (BioTek Industries). Statistical analyses were performed on log-transformed data.

### Antigen specific Isotype ELISA

MaxiSorp plates (Nunc) were coated with 50 μl of 2 μg/mL or 50ul of 5 μg/mL ng/well SARS-CoV-2 full-length spike (FL-S) protein overnight at 4 °C for detection of IgG (250ng/well) or IgM and IgA (500ng/well), respectively, prior to washing in PBS/Tween (0.05% v/v) and blocking with Blocker Casein in PBS (Thermo Fisher Scientific) for 1 h at room temperature (RT). Standard positive serum (pool of mouse serum with high endpoint titre against FL-S protein), individual mouse serum samples, negative and an internal control (diluted in casein) were incubated for 2 hrs at room temperature for detection of specific IgG or 1h at 37 °C for detection of specific IgM or IgA. Following washing, bound antibodies were detected by addition of alkaline phosphatase (AP)-conjugated goat anti-mouse IgG (Sigma-Aldrich) or anti-mouse IgM (Abcam) or anti-mouse IgA (SouthernBiotech) for 1h at room temperature and addition of p-Nitrophenyl Phosphate, Disodium Salt substrate (Sigma-Aldrich). An arbitrary number of ELISA units (EU) were assigned to the reference pool and optical density values of each dilution were fitted to a 4-parameter logistic curve using SOFTmax PRO software. ELISA units were calculated for each sample using the optical density values of the sample and the parameters of the standard curve. IgM limit of detection was defined a 2 ELISA Units, IgA limit of detection set as 6 ELISA Units. All data was log-transformed for statistical analyses.

### Antigen Specific IgG Subclass ELISAs

MaxiSorp plates (Nunc) were coated with 50 μL of 2 μg/mL per well of SARS-CoV-2 FL-S protein overnight at 4 °C prior to washing in PBS/Tween (0.05% v/v) and blocking with Blocker Casein in PBS (Thermo Fisher Scientific) for 1 h at room temperature (RT). For detection of IgG subclasses all serum samples were diluted to 1 total IgG EU and incubated at 37 °C for 1 h prior to detection with Alkaline Phosphatase conjugated anti-mouse IgG subclass-specific secondary antibodies (Southern Biotech or Abcam) incubated for 1 h at 37 °C. The results of the IgG subclass ELISA are presented using optical density values.

### Avidity ELISA

Anti-SARS-CoV-2 spike-specific total IgG antibody avidity was assessed by sodium thiocyanate (NaSCN)-displacement ELISA. Nunc MaxiSorp ELISA plates (Thermo Fisher Scientific) coated overnight at 4⍰°C with 2μg/well SARS-CoV-2 FL-S protein diluted in PBS were washed with PBS/Tween (0.05% v/v) and blocked for 1⍰h with 100⍰μl per well of Blocker Casein in PBS (Thermo Fisher Scientific) at 20⍰°C. Test samples and a positive control serum pool were diluted in blocking buffer to 1 total IgG EU and incubated for 2⍰h at 20⍰°C. After washing, increasing concentrations of NaSCN (Sigma-Aldrich) diluted in PBS were added and incubated for 15⍰min at 201°C. Following another wash, bound antibodies were detected by addition of AP-conjugated goat anti-mouse IgG (Sigma-Aldrich) for 1 h at room temperature and addition of p-Nitrophenyl Phosphate, Disodium Salt substrate (Sigma-Aldrich). For each sample, concentration of NaSCN required to reduce the OD405 to 50% of that without NaSCN (IC50) was interpolated from this function and reported as a measure of avidity.

### Antigen specific B cell staining

Spike, RBD and a decoy NANP_9_C (repeat region from *P. falciparum* CSP protein) tetramers were prepared in house by mixing biotinylated proteins with streptavidin conjugated flurochromes (A488, A647 or r-PE) in a 4:1 molar ratio and incubating on ice for 30 minutes. Splenocytes were stained with Spike-PE and RBD-A647 at a final concentration of 0.04mM, whilst decoy tetramer, NANP_9_C-A488 was used at a final concentration of 0.4mM. Splenocytes were stained with Live-Dead Aqua and Fc block (anti-CD16/32 mAb, Clone 93) prior to staining with the antibody cocktail containing NANP_9_C-Alexa488, GL7-PerCPCy5.5 (Clone GL7), CD138 BV421 (Clone 281-2), CD95-BV605 (Clone SA367H8), CD4-BV650 (Clone GK1.5), CD279-BV711 (Clone 29F.1A12), CD19-BV780 (6D5), RBD-A647, IgD-A700 (Clone 11-26c.2a), IgM-APCCy7 (Clone 11/41), Spike-PE, CD38-PECY5 (Clone 90), CD69 PeCy7 (Clone H1.2F3), CD45R-BUV395 (Clone RA3-6B2) and CD3-BUV496 (Clone 145-2C11), antibodies purchased from BioLegend, BD or Invitrogen. Antigen specific B cells were identified by gating on LIVE/DEAD negative, size (FSC-A vs SSC), doublet negative (FSC-H vs FSC-A), CD45RA^+^, CD19^+^ and NANP-A488^−^, RBD-A647^+^ and Spike-PE^+^ followed by staining for germinal centre or class switched B cells (Figure S2A). The total number of cells was calculated by multiplying the frequency of each population, expressed as a percentage of total lymphocytes, by the total number of lymphocytes counted for each individual mouse spleen sample.

### ELISpot and ICS staining

Spleen single cell suspension were prepared by passing cells through 70μM cell strainers and ACK lysis prior to resuspension in complete media. Splenocytes were stimulated 15mer peptides (overlapping by 11) spanning the length of SARS CoV2 protein, with peptide pools subdivided into peptides spanning the S1 and S2 region of spike (Table S1). For analysis of IFNγ production by ELISpot, splenocytes were stimulated with two pools of S1 peptides (pools 1 and 2) and two pools of S2 peptides (pools 3 and 4) (final concentration of 2μg/mL) on IPVH-membrane plates (Millipore) coated with 5μg/mL anti-mouse IFNγ (AN18). After 18-20 hours of stimulation at 37 °C, IFNγ spot forming cells (SFC) were detected by staining membranes with anti-mouse IFNγ biotin (1mg/mL) (R46A2) followed by streptavidin-Alkaline Phosphatase (1mg/mL) and development with AP conjugate substrate kit (BioRad, UK).

For analysis of intracellular cytokine production, cells were stimulated at 37 °C for 6 hours with 2μg/mL a pool of S1 (ELISpot pools 1 and 2) or S2 (ELISpot pools 3 and 4) peptides (Table S1), media or positive control cell stimulation cocktail (containing PMA-Ionomycin, BioLegend), together with 1μg/mL Golgi-plug (BD) and 2μl/mL CD107a-Alexa647 (Clone 1D4B). Following surface staining with CD3-A700 (Clone 17A2), CD4-BUV496 (Clone GK1.5), CD8-BUV496 (Clone 53-6.7), CD44-BV780 (Clone IM7), CD62L-BV711 (Clone MEL-14), CD69-PECy7 (Clone H1.2F3) and CD127-BV650 (Clone A7R34) or CD127-APCCy7 (Clone A7R34), cells were fixed with 4% paraformaldehyde and stained intracellularly with TNFa-A488 (Clone MP6-XT22), IL2-PerCPCy5.5 (Clone JES6-5H4), IL4-BV605 (Clone 11B11), IL10-PE (Clone JES5-16E3) and IFNγ-e450 (Clone XMG1.2) diluted in Perm-Wash buffer (BD). Sample acquisition was performed on a Fortessa (BD) and data analyzed in FlowJo V10 (TreeStar). An acquisition threshold was set at a minimum of 5000 events in the live CD3^+^ gate. Antigen specific T cells were identified by gating on LIVE/DEAD negative, doublet negative (FSC-H vs FSC-A), size (FSC-A vs SSC), CD3^+^, CD4^+^ or CD8^+^ cells and each individual cytokine or “cytokine positive” comprised of a combination of “CD107a or IFNγ or TNFα or IL2 or IL4 or IL10) populations. Cytokine positive responses are presented after subtraction of the background response detected in the corresponding media stimulated control sample for each mouse and summing together the response detected to each pool of peptides. T effector (Teff) cells were defined as CD62L^low^ CD127^low^, T effector memory (Tem) cells defined as CD62L^low^ CD127^hi^ and T central memory (Tcm) cells defined as CD62L^hi^ CD127^hi^ (Figure S3B). The total number of cells was calculated by frequency of the background corrected population (expressed as a percentage of total lymphocytes) by the total number of lymphocytes counted in each individual spleen sample.

### Statistical analysis

All graphs and statistical analysis were performed using Prism v9 (Graphpad). For analysis of vaccination regimen against a single variable (eg IgG level), data was analysed with a one-way anova (Kruskal-Wallis) followed by post-hoc positive test. For analysis of vaccination regimen against multiple variables (eg each individual cytokine or T cell subset) the data was analysed with a two-way analysis of variance, where a significant difference was observed, a post-hoc analysis was performed to compare the overall effect of vaccination regimen. In graphs where a significant difference was observed between multiple vaccine groups, the highest p value is displayed on the graph. All data displayed on a logarithmic scale was log_10_ transformed prior to statistical analysis (ELISA Units, Neutralisation Titres, Total Cell Numbers).

## Acknowledgments

The authors would like to thank D. Pulido for provision of spike and RBD proteins, BMS staff for animal husbandry and management and A. Worth, J.Furze, M. Mykhaylyk and R. Evans for facilities support.

## Author Contributions

AJS, MU, SB-J, CB, KS, KM & PMK performed experiments. AJS, MU, HS, CG, EA & AT performed animal procedures and/or sample processing. AJS, SBJ, KH & PMK, analyzed data. AJS, TL, PMK, RS & SG designed the study. AJS, SBJ, TB wrote the manuscript. All authors reviewed the final version of the manuscript.

## Figure legends

**Fig S1:**
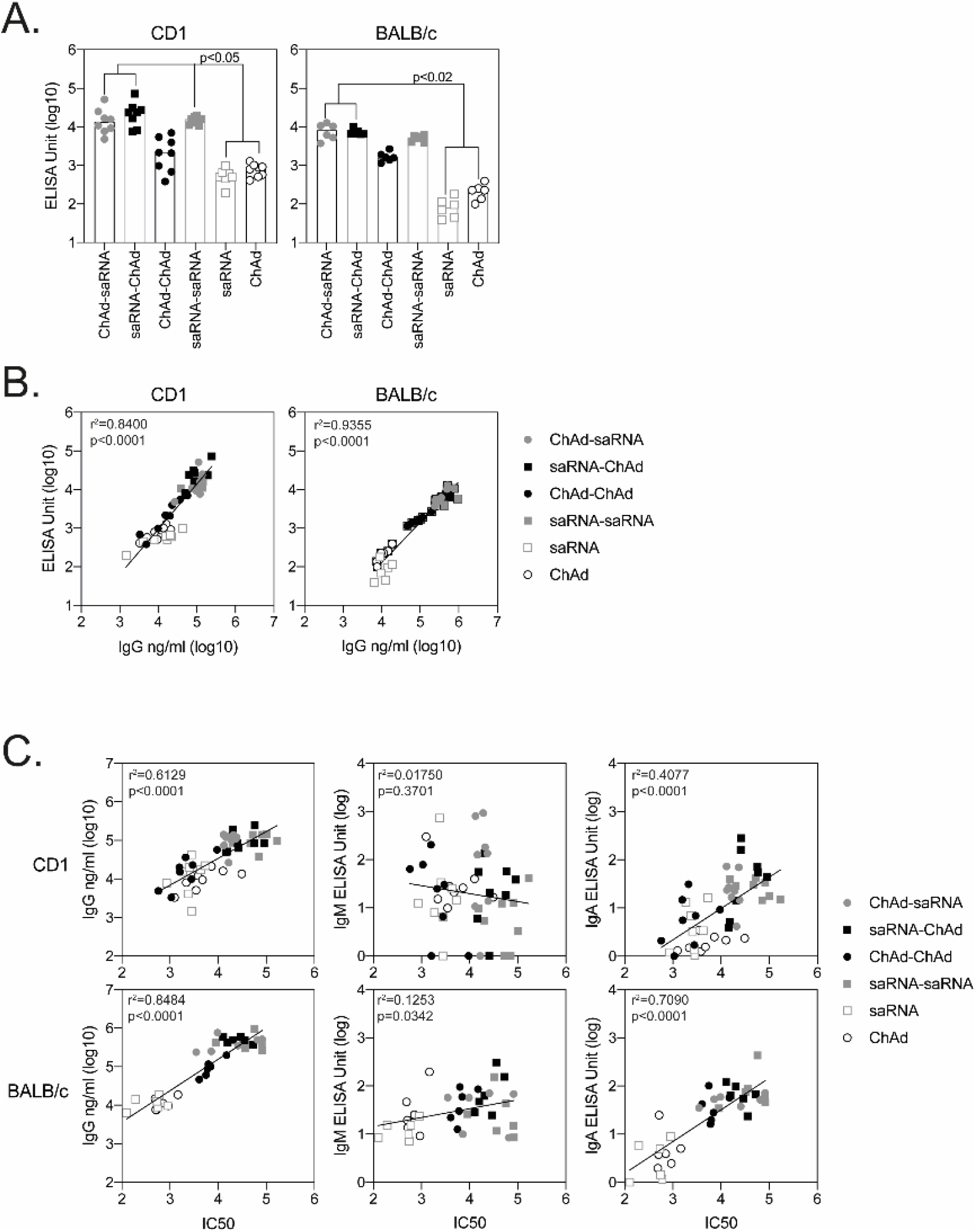
Comparison of assays measuring antibody responses after vaccination. SARS-CoV-2 spike-specific IgG responses were measured in a standardised ELISA **(A.)** and compared to IgG responses measured as a concentration **(B.) C.)** Graphs show relationship between IgG, IgM and IgA responses to SARS-CoV-2 pseudoneutralisation IC50 values presented by linear regression on log-transformed values. Data points represent individual animals from all groups together, r^2^ and p values are indicated on each graph.

**Fig S2:**
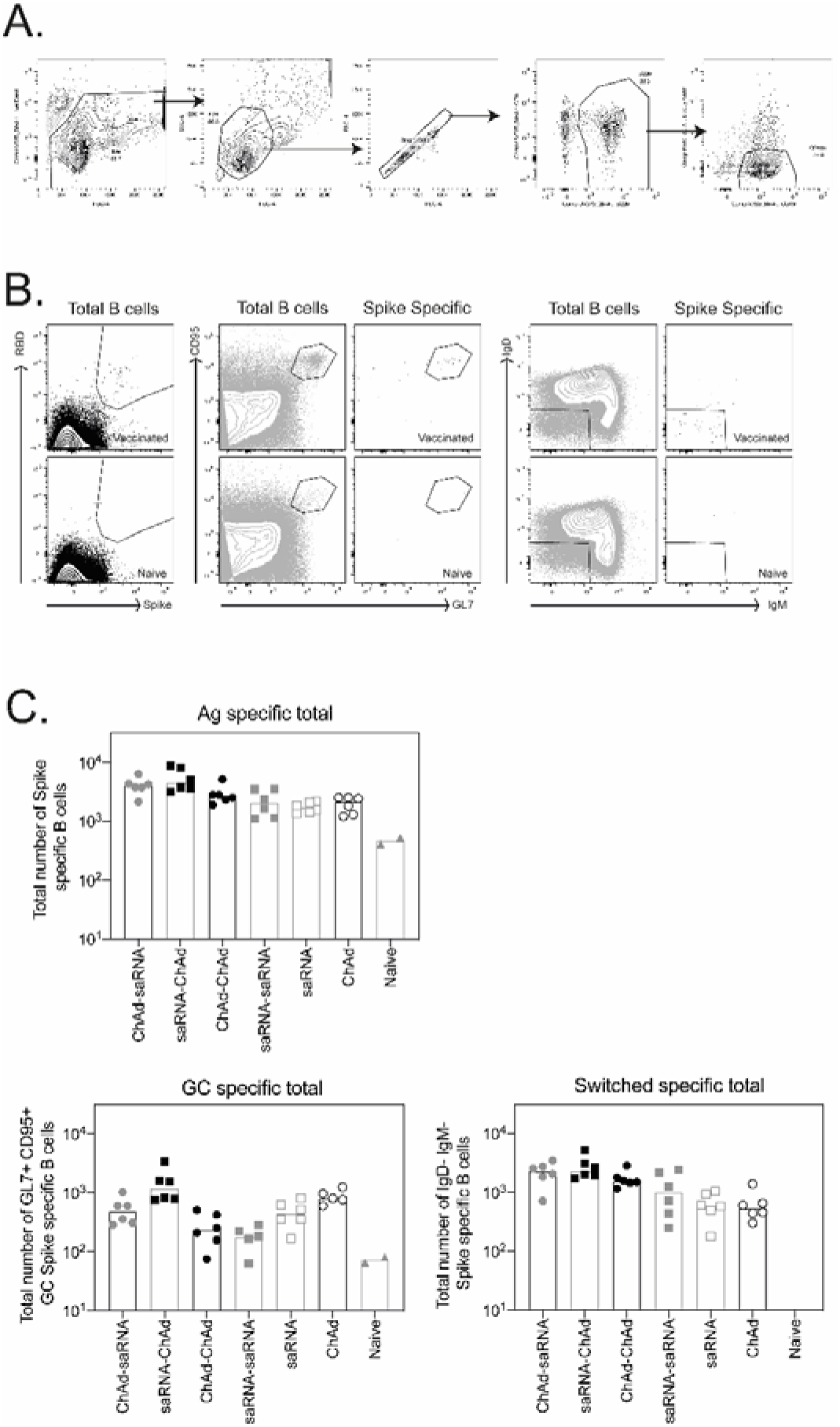
Antigen specific B cell responses. SARS-CoV-2 spike-specific B cells responses were measured in the spleen of BALB/c mice 3 weeks after the final immunisation. **A.)** Plots show the gating strategy for identification of B cells. **B.)** antigen specific cells were identified by positive binding to spike-PE and RBD-A647 tetramers with an antigen specific population visible in vaccinated animals (top plot) but not naïve mice (lower panel). Germinal center (GC) B cells were identified as GL7^+^ and CD95^+^, switched B cells identified as IgD^−^ and IgM^−^ antigen-specific B cells. C.) Graphs show the total number of antigen-specific B cells, antigen-specific GC B cells and antigen-specific switched B cells, data points are representative of individual mice, median responses per group indicated by bars.

**Fig S3:**
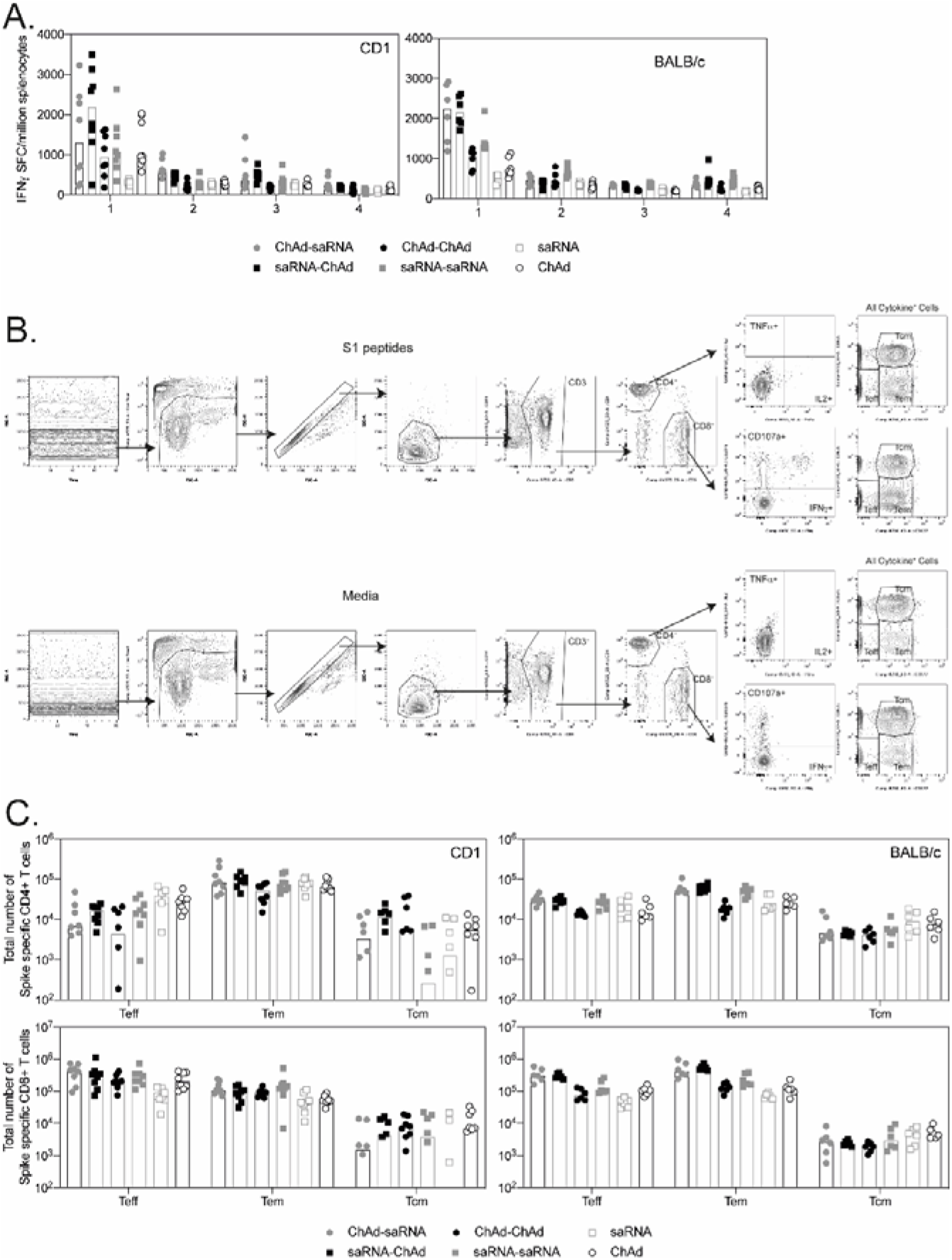
T cell responses measured by ELISpot and ICS. **A.)** Graphs show the IFNγ SFC detected to each pool of peptides by ELISpot. Each data point represents an individual mouse, bars represent the median response per group. **B.)** Plots show the gating strategy used to identify antigen specific T cell responses. **C.)** Graphs show the total number of Teff, Tem and Tcm responses detected in the spleens of mice. Each point indicates an individual mouse, bars represent the median response per group.

